# An ATF4-mediated transcriptional adaptation of electron transport chain disturbance primes progenitor cells for proliferation *in vivo*

**DOI:** 10.1101/425744

**Authors:** Sebastian Sorge, Jonas Theelke, Christian Altbürger, Ingrid Lohmann

**Author notes:** Author for correspondence (I.L.). Mailing address of corresponding author, Ingrid Lohmann, University of Heidelberg, Im Neuenheimer Feld 230, D-69120 Heidelberg, FX, Germany, EM.

## Abstract

The mitochondrial electron transport chain (ETC) enables many important metabolic reactions, like ATP generation and redox balance. While the vital importance of mitochondrial function is obvious, the cellular response to defects in mitochondria and in particular the modulation of signalling pathway outputs is not understood. Using the *Drosophila* eye as model, we show that the combination of Notch signalling and a mild attenuation of the ETC via knock-down of COX7a causes massive cellular over-proliferation. The tumour like growth is caused by a transcriptional response through the eIF2α-kinase PERK and ATF4, a stress-induced transcription factor, which activates the expression of many metabolic enzymes, nutrient transporters and mitochondrial chaperones. We find this stress adaptation to be beneficial for progenitor cell fitness upon ETC attenuation. Activation of the ATF4 mediated stress response renders cells sensitive to proliferation induced by the growth-promoting Notch or Ras signalling pathways, leading to severe tissue over-growth. In sum, our results suggest ETC function is monitored by the PERK-ATF4 pathway, a cellular adaptation hijacked by growth-promoting signalling pathways in situations of oncogenic pathway activity.

## INTRODUCTION

Controlling cell proliferation is one of the major challenges of multicellular life, both during phases of growth in developing organisms and phases of homeostatic cell-replenishment essential in adult animals. Lack of appropriate control can lead to severe disorders at any stage of life, including cancer. A century ago, Otto Warburg discovered a metabolic shift of cancer cells from respiration towards aerobic glycolysis and hypothesised that defective mitochondrial respiration triggers cancer development [1]. While mitochondria are not dysfunctional in many cancers, the switch to a glycolytic metabolism in cancer cells has been confirmed as a general feature of most cancers [2]. In addition, it has become clear more recently that proliferating mammalian cells generally rely on a glycolytic metabolism, presumably because high glycolytic flux allows for intermediates to be used for anabolic reactions, such as nucleotide biosynthesis [3]. Besides these intriguing findings, most of today’s knowledge of how organisms control proliferation stems from the study of signalling pathways, which are frequently mutated in cancer (oncogenes and tumour suppressors) and causally linked to the uncontrolled growth of tumours [4]. In recent years, several studies have implicated a direct metabolic regulation of cell cycle progression [5,6] and indirect effects on proliferation through reactive oxygen species (ROS) released by a dysfunctional mitochondrial electron transport chain [6,7]. In addition, it was shown that the nutritional status of imaginal cells, proliferating epithelial tissues in *Drosophila* that give rise to most external structures of the adult fly during metamorphosis, directly impacted on their proliferative behaviour [8]. Together, these studies highlight the importance of intersections between classical signalling pathways and metabolism in proliferating imaginal progenitors *in vivo*. Yet, most clonal assays focus on non-cell autonomous effects, omitting direct consequences of metabolic defects or adaptations on the physiology of (imaginal) progenitor cells and development of the tissues.

Here, we used a tissue-specific GAL4-driver based labelling and manipulation approach targeting all eye progenitors to elucidate the interplay of metabolism and proliferation control. Compared to clonal analysis, this setup allowed us to study cell-autonomous proliferation effects independently of competition between cells of different fitness, a process dominant in most clonal analyses. We found that eye progenitors autonomously activate a transcriptional stress response upon impairment of the mitochondrial electron transport chain, mediated through PERK and ATF4. We further demonstrated that this adaptation, which helps eye progenitors to follow their developmental program could be hijacked by growth-promoting pathways like Notch or Ras to enhance their proliferative effects. Our results suggest that ATF4-mediated transcriptional adaptation provides a cell-autonomous response to ETC defects, altering cellular behaviour through metabolic adaptation.

## RESULTS

### Mild reduction of COX7a enhances Notch-induced proliferation

In order to identify new regulators of cell proliferation, we used RNA-seq expression data of a previously established eye tumour model in *Drosophila*, in which loss of the transcription factor Cut was shown to cooperate with Notch signalling to induce tumorigenesis [9] and tested down-regulated genes for tumour suppressor-like activity. To this end, we reduced their expression in the *Drosophila* eye by RNAi using the *eyeless* (*ey*)-GAL4 driver and screened for their ability to modify the well-described Notch-mediated mild over-proliferation induced by Delta over-expression (Dl^OE^) [10] (Figures 1A, 1D and S1A). Among the 109 genes tested (Supplementary Table 1) we found one, the mitochondrial respiratory chain subunit *Cytochrome C*-*oxidase subunit 7a* (*COX7a*), which enhanced the slight over-proliferation induced by Dl^OE^ when its expression was reduced. In this context, we recovered over-proliferating eye imaginal discs (Figures 1D’, 1E’ and S1B), leading to strongly folded adult eyes (Figures 1D, 1E and S1A), resembling tumorous tissue overgrowth. Interestingly, raising GAL4 activity using higher temperature (29°C instead of 25°C) increased overgrowth of Dl^OE^, COX7a^RNAi^ larval eye discs (Figures 1E’ and 1F’), but also affected cell differentiation, as we observed a marked loss of ELAV-positive cells in the ventral domain of eye imaginal discs (Figures 1E’ and 1F’), and reduced and malformed eyes in adult flies (Figures 1E and 1F). Strikingly, RNAi based interference with *COX7a* alone, which had been shown to cause general developmental defects and lethality due to attenuation of Complex 4 activity [11], negatively impacted on proliferation, as *COX7a* depleted *Drosophila* S2R+ cells (Figure S1D) were unable to divide (Figure S1C), and consistently the sizes of eye discs (Figures 1A’ and 1B’) and of adult eyes (Figures 1A, 1B and S1A) were reduced. Enhancement of RNAi further reduced eye size and also affected differentiation in COX7a^RNAi^ eyes (Figures 1C and 1C’). Co-expression of the apoptosis inhibitor p35 [12], did not rescue eye size (Figures 1G-1I), excluding enhanced apoptosis induction as the primary cause of the smaller eye size phenotype in COX7a^RNAi^ or over-proliferation of Dl^OE^, COX7a^RNAi^ animals. These results suggested that increased loss of *COX7a* in combination with Notch activation triggered over-proliferation in eye progenitors, and additionally impaired either their capacity to enter the differentiation program or induced a switch towards a different (cuticle) fate.

**Figure 1:**
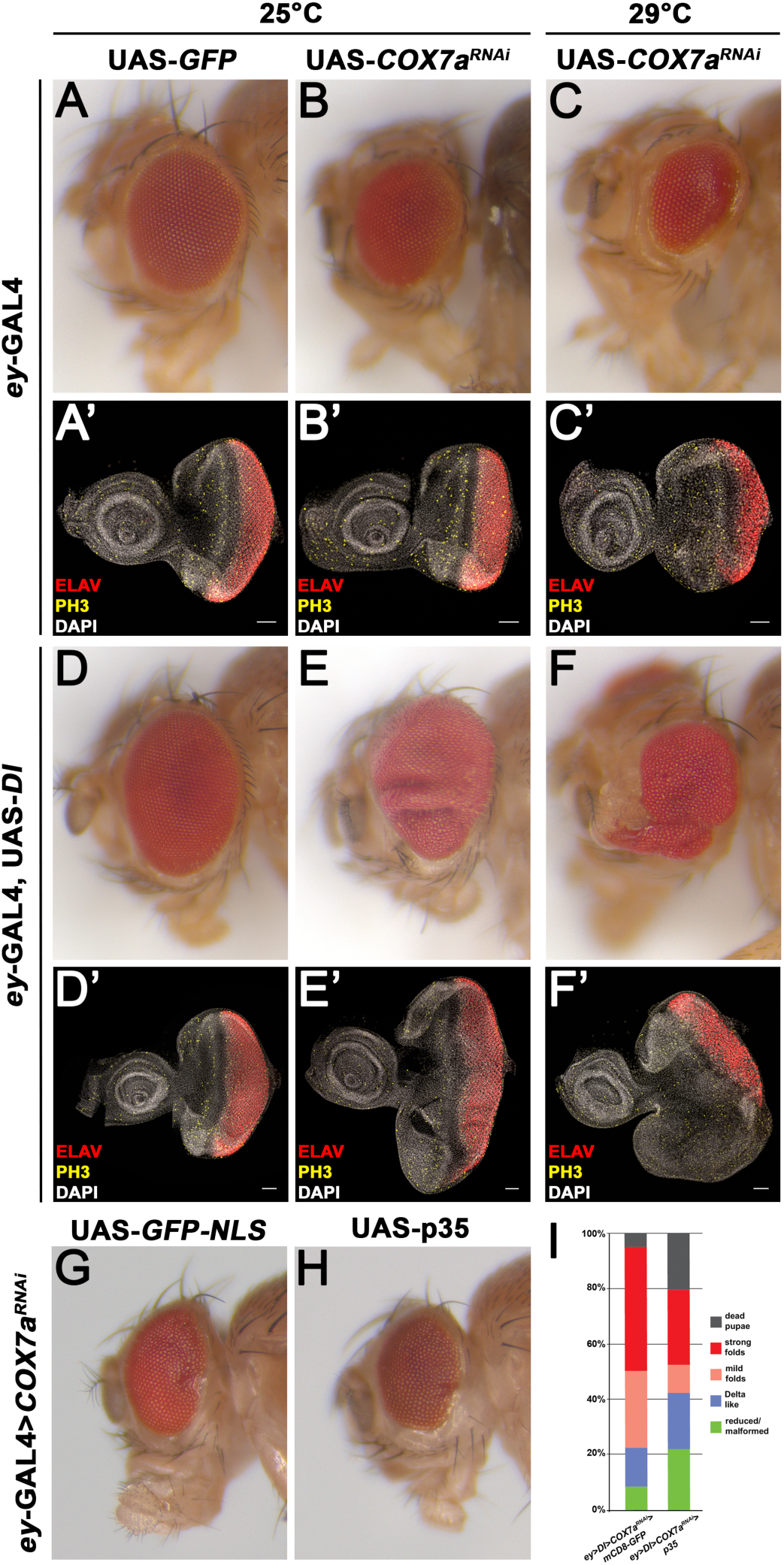
Reduction of COX7a enhances Notch-induced proliferation. **A-F** Adult eyes of the indicated genotypes. Animals were reared at 25°C (A, B, D, E) or 29°C (C, F) on standard fly food. COX7a knock-down reduced adult eye size in a dose-dependent manner (compare B, C and control A). Eyes of Delta over-expressing animals (D) were elongated along the dorso-ventral axis, due to increased proliferation of the ventral compartment. Knock-down of COX7a in the Delta background (E) increased eye size and led to folding of the flat epithelium. Adult survivors at 29°C exhibited small, malformed eyes and head capsule (F). **A’-F’** Sum-projections of larval late L3 eye-imaginal discs of the indicated genotypes. Eye discs are stained for ELAV (red, labelling photoreceptors), phospho-Histone 3 (PH3) (yellow, labelling mitotic cells) and DAPI (white, labelling nuclei). Animals were reared at 25°C (A’, B’, D’, E’) or 29°C (C’, F’) on standard fly food. At 29°C (C’), dividing cells in the first mitotic wave (anterior to the morphogenetic furrow) and second mitotic wave (posterior o the morphogenetic furrow) appeared reduced. Delta over-expressing discs (D’) show an enlarged ventral compartment. Knock-down of COX7a in the Delta background (E’) substantially increased size of the epithelium and showed a high density of proliferating cells in the ventral domain, where the progression of differentiation (ELAV, red) appeared to be slowed down. At 29°C, heavily overgrown larval discs were recovered (F’), showing strong defects of cells to enter differentiation in the ventral compartment. **G, H** Adult eyes from offspring of crosses of *ey*>*COX7a^RNAi^* females with *UAS*-*GFP^NLS^* males (G) as control or UAS-*p35* males (H) to inhibit apoptosis. Eye size was reduced in the absence of apoptosis, showing that cell death itself is not the reason for the small eye phenotype. Larvae were reared at 25°C under nutrient restriction. **I** Quantification of adult eye phenotypes from offspring of crosses of *ey*>*Delta*>*COX7a^RNAi^* females with UAS-*mCD8*-*GFP* or UAS-*p35* males. Inhibition of apoptosis through p35 expression did not inhibit over-proliferation. Larvae were reared at 25°C under nutrient restriction. Anterior is to the left, dorsal is up, scale bar represents 50μm in all microscope images. See also Fig S1.

In sum, our data showed that a mild disturbance of the ETC induced by *COX7a* knock-down modulated the Notch signalling pathway, thereby inducing the over-proliferation of an epithelial tissue system, while stronger interference with *COX7a* affected in addition the differentiation of progenitor cells. As we were interested in understanding the modification of proliferation (but not of differentiation) by changes in ETC activity, we focused our analysis on the mild knock-down of *COX7a* at 25°C.

### Notch and COX7a induce independent transcriptional responses that cooperate in proliferation control

The Notch pathway affects cell behaviour primarily by controlling the transcriptional output via the processed Notch^intra^ domain interacting with the Su(H) transcription factor [13]. We assumed the COX7a mediated enhancement of over-proliferation in Dl^OE^, COX7a^RNAi^ animals to be caused by a modification of the Notch induced transcriptional response. Thus, we globally profiled transcriptome changes by performing microarray analysis of early L3 eye-antennal discs that were at the onset of differentiation. This analysis confirmed activation of generic and eye-specific Notch target genes, including *Enhancer of split* complex genes like *E(spl)m2*, *E(spl)m3* and *E(spl)m7* and *eyegone* (*eyg*), when the Notch pathway was activated by Dl^OE^ (Figure 2A), and showed that the tissue was not compromised by Delta over-expression, as stress or other pathway genes were not induced (Figures 2C, S2D, S2F). Interference with COX7a induced a completely different transcriptional response, which hardly overlapped with the changes caused by Dl^OE^ (Figures 2F and 2G). We found transcripts for several glycolytic enzymes including lactate dehydrogenase (LDH) (Figure 2D), amino acid and sugar transporters (Figure 2B), as well as mitochondrial chaperones (Figure 2C) increased in COX7a^RNAi^ but not in Dl^OE^ imaginal discs. Using a *LDH*-*GFP* enhancer trap [14] we confirmed LDH expression in L3 eye-antennal discs upon *COX7a* knockdown (either alone or in combination with Dl^OE^) (Figure 2E and 2H). These results suggested that eye progenitors responded to Delta over-expression by over-activating the Notch pathway and mounted a specific stress-response upon knockdown of COX7a, without activating typical stress pathways like Nrf2 or JNK (Figures S2D - S2F). However in contrast to our expectations, we did not observe enhanced induction of COX7a^RNAi^ or Dl^OE^ targets in Dl^OE^, COX7a^RNAi^ eye discs (Figures 2A-2D), nor did we observe the induction of new targets (besides a few) in COX7a^RNAi^, Dl^OE^ eye discs (Figure S2G). These results demonstrated that the COX7a^RNAi^, Dl^OE^ induced over-proliferation was not caused by the transcriptional cooperation of Notch and an unknown COX7a nuclear effector but that these two programs act independently.

**Figure 2:**
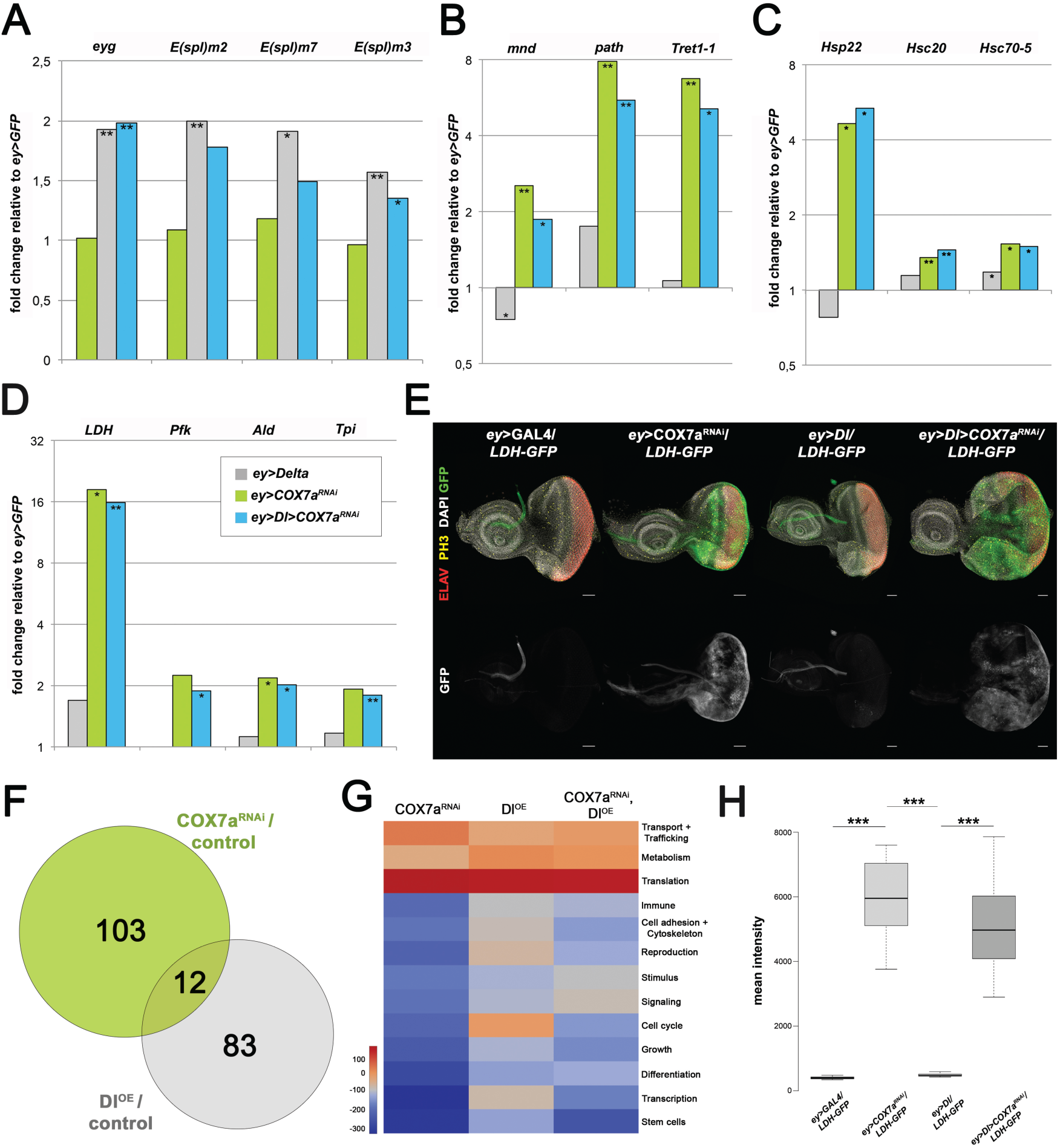
ETC impairment and Notch over-activation cause different nuclear responses. **A-D** Fold change of selected target genes of the Notch pathway (A), nutrient transporters (B), mitochondrial chaperones (C) and glycolytic enzymes (D) shown in the respective genotypes relative to *ey>GFP* control. Asterisks indicate significance of fold-change to *ey*>*GFP* (^*^ = p<0,05; ^**^ = p<0,01). **E** Sum-projection of late L3 eye-antennal discs stained for ELAV (red, labelling photoreceptor clusters), phospho-Histone 3 (PH3) (yellow, labelling mitotic cells), DAPI (white, labelling nuclei) and carrying the *LDH*-*GFP* (green) enhancer trap. Larvae were reared at 25°C on standard fly food. Control (*ey*-*GAL4*) or *DI^OE^* discs did not express any detectable GFP. *COX7a^RNAi^* induced GFP transcription primarily in progenitor cells. **F** Venn diagram showing that *COX7a^RNAi^* and *DI^OE^* induced primarily distinct sets of target genes (fold>1,5; p<0,05). **G** Heat-map displaying presence of genes belonging to higher-order categories in genetic backgrounds. The different representation of GO term categories reflects the different transcriptional responses induced by *COX7a^RNAi^* and *DI^OE^*. The colour range corresponds to the delta rank of genes annotated to the category that also appear in the sample: red colour represents enrichment of higher-expressed genes, blue colours categories enriched with lower-expressed genes. Rows are hierarchically clustered using Euclidean distance with complete linkage. **H** Quantification of LDH-GFP signal of larval discs as shown in E (n=6). Asterisks indicate p-value according to unpaired t-test: ^*^ p<0,05; ^**^ p<0,01; ^***^ p<0,001. Anterior is to the left, dorsal is up, scale bar represents 50μm in all microscope images. See also Fig S2.

We noticed however that food quality altered COX7a^RNAi^ induced phenotypes, as diets with reduced protein content enhanced over-proliferation of eye tissue in Dl^OE^, COX7a^RNAi^ animals (Figures S2B and S2C) and promoted eye size reduction in COX7a^RNAi^ animals (Figure S2B), while over-proliferation of adult eyes in Dl^OE^, COX7a^RNAi^ animals was absent on a pure yeast (high protein) diet (Figure S2B). And consistent with the enhancement of the phenotypic strength under nutrient restriction, many more targets were induced by COX7a^RNAi^ when animals were raised under poor diet conditions (472 vs. 115), while the majority of targets induced on normal food were confirmed (86/115) (Figure S2A). These results suggested that amino acid limitation directly altered eye progenitor cells response to COX7a knockdown through modulation of intracellular signalling or alternatively a dependence on systemic hormone signalling.

Together, these results revealed that COX7a knock-down induced a transcriptional response independent of Notch via an unknown nuclear effector activating genes with diverse cellular functions.

### ATF4 mediates the nuclear response induced by ETC disturbance

In order to uncover the pathway mediating the nuclear response induced by COX7a knock-down, we genetically screened for candidates able to modify the Dl^OE^, COX7a^RNAi^ over-proliferation phenotype. The induction of mitochondrial chaperones indicated that COX7a^RNAi^ cells were compromised and stressed, thus we tested many of the well-known stress-response or proliferation-inducing pathways, including JNK, Wnt, AMPK, Nrf2, and HIF1α. Although we observed a slight activation of the JNK pathway as evidenced by reporter lines and target genes (Figures S2D and S2E), none of these pathways was able to rescue Dl^OE^, COX7a^RNAi^-induced over-proliferation when their activity was reduced (data not shown). Thus, we excluded them as mediators of the combinatorial COX7a and Notch induced effects. Another stress-induced transcription factor adapting cells to various insults such as ER protein folding stress is Activating Transcription Factor 4 (ATF4; *cryptocephal* in *Drosophila*). Consistent with a possible involvement of ATF4, we identified the human ATF4 motif as the highest-ranking motif among the regulatory sequences of COX7a^RNAi^-induced genes (22/114; Figure 3A) using iRegulon prediction of motif enrichment [15]. Interestingly, in *Drosophila* S2R^+^ cells, ATF4 was shown to induce LDH and glycolytic enzymes upon ER stress [16], which we also found activated in eye discs (Figures 2D, 2E and 2H), and in mammals, ATF4 regulates various amino acid transporters homologous to the ones we recovered in our *in vivo* transcriptome profiling [17]. In order to test a potential involvement of ATF4, we performed knock-down experiments in Dl^OE^, COX7a^RNAi^ animals and found that down-regulation of ATF4 completely rescued the over-proliferation in larval discs and adults (Figures 3C, 3E and 3G), resembling phenotypically Dl^OE^ eyes (Figure 1D). Using an antibody against ATF4 [18] we detected transient ATF4 expression upon COX7a^RNAi^ in early L3 eye progenitors anterior to the morphogenetic furrow, while control and late L3 eye discs lacked detectable ATF4 protein expression (Figures 3C and S3A). Importantly, the activation of the vast majority of COX7a^RNAi^ targets (431/472) required ATF4 function, as these targets were not induced in COX7a^RNAi^, ATF4^RNAi^ discs (Figure S3D). Likewise, the majority of target genes induced by the combination of Dl^OE^ and COX7a^RNAi^ required ATF4 (Figure 3B). These target genes include all COX7a^RNAi^-induced genes shown in Figures 2B-2D, both in the presence (Figure 3D) or absence (Figure S3C) of Dl^OE^. The LDH-GFP reporter acted as a bona fide reporter for ATF4 activity, as GFP expression was undetectable in COX7a^RNAi^, ATF4^RNAi^ eye discs (Figure 3F).

**Figure 3:**
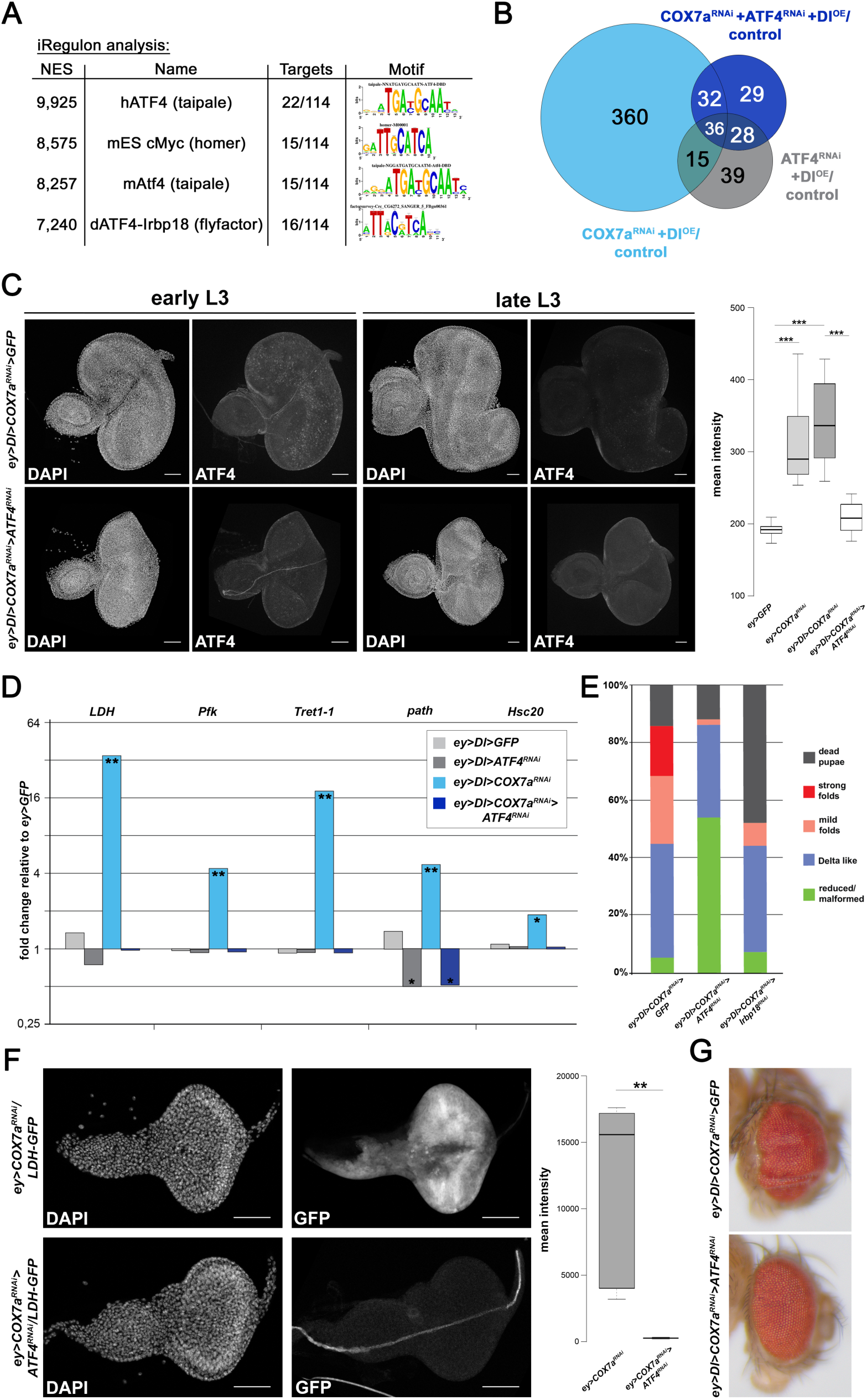
PERK-ATF4 mediate nuclear response of ETC impairment. **A** Highest ranking transcription factor motifs according to iRegulon analysis under standard settings (9713 PWMs; 5kb upstream, 5’-UTR and 1^st^ intron with standard cut-offs). The human ATF4 motif was the highest-ranking motif, followed by a cMyc motif (that is a near-identical reverse-complement of the hATF4 motif), mouse Atf4 and a *Drosophila* heterodimer of ATF4 and Irbp18. **B** Venn diagram depicting dependency of *Dl^OE^, COX7a^RNAi^* induced target genes on *ATF4* function. For this microarray analysis, larvae were reared at 25°C on nutrient restriction food. The majority of *Dl^OE^*, *COX7a^RNAi^* induced target genes (360/443) were not induced in *DI^OE^*, *COX7a^RNAi^*, *ATF4^RNAi^* discs. **C** Sum-projection of early or late L3 eye-antennal discs stained for DAPI (labelling nuclei) and ATF4. Larvae were reared at 25°C on nutrient restriction food. ATF4 was detected in early L3 progenitors, but not in late L3 discs. *ATF4^RNAi^* abolished ATF4 protein induction and over-proliferation. Quantification (to the right) of ATF4 signal in early L3 eye discs of these genotypes and the ones of Figure S3A. Asterisks indicate p-value according to unpaired t-test: ^*^ p<0,05; ^**^ p<0,01; ^***^ p<0,001. **D** Fold change of selected *COX7a^RNAi^*-induced genes in the *Delta^OE^* background (relative to *ey*>*GFP* control) according to second microarray analysis under nutrient restriction. Induction of these genes in response to *COX7a^RNAi^* requires *ATF4* function. Asterisks indicate significance of fold-change to *ey*>*GFP* (^*^ = p<0,05; ^**^ = p<0,01). **E** Quantification of adult eye phenotypes in crosses of *ey*>*Df^OE^,COX7a^RNAi^* females with either UAS-GFP (control), *ATF4^RNAi^* or *Irbp18^RNAi^* males. Larvae were reared at 25°C on nutrient restriction food. Knockdown of *ATF4* and its cofactor *Irbp18* repressed over-proliferation. **F** Sum-projection and quantification of LDH-GFP signal in early L3 eye-antenna discs. In the absence of *ATF4* function, LDH-GFP is not detectable (as in controls). Quantification (to the right) of GFP signal in early L3 eye discs of these genotypes. Asterisks indicate p-value according to unpaired t-test: ^*^ p<0,05; ^**^ p<0,01; ^***^ p<0,001. **G** Representative adult eyes of *ey*>*Dl^OE^*,*COX7a^RNAi^*, *GFP* and *ey*>*Dl^OE^*,*COX7a^RNAi^*, *ATF4^RNAi^* flies, showing that *COX7a^RNAi^*-mediated over-proliferation requires *ATF4*. Anterior is to the left, dorsal is up, scale bar represents 50μm in all microscope images. See also Figure S3.

We next sought to understand which of the ATF4 targets enhanced proliferation and screened several candidates in the Dl^OE^, COX7a^RNAi^ background. Amongst the COX7a^KD^-induced genes we identified a single gene, called *Irbp18*, whose knockdown rescued over-proliferation, qualitatively and quantitatively similar to ATF4^RNAi^ (Figure 3E). *Irbp18* encodes the homologue of C/EBPγ and had been shown to be one of two heterodimeric binding partners for ATF4 [19], which was shown before to regulate most ATF4-dependent target genes of the integrated stress response (ISR) together with ATF4 in mouse cells [20]. Screening of genes with direct metabolic functions did not reveal a single gene that was similarly required for over-proliferation as the upstream regulators. At best, we found that the knockdown of *pathetic* (*path*), a gene encoding an amino acid transporter or sensor [21], and the *Drosophila* homolog of BCAT1, an branched-chain amino acid biosynthetic enzyme, slightly attenuated over-proliferation (Figure S3E). Thus, we assume the induction of a complex network of ISR genes to be required to enhance Notch-mediated over-proliferation.

Taken together, these results demonstrated that ATF4 and C/EBPγ acted as the main transcriptional effectors downstream of COX7a-mediated mild ETC disturbance and suggested that multiple ATF4 and Notch induced target genes cooperate in eye progenitor over-proliferation.

### PERK mediates ATF4 activation upon ETC disturbance

In a next step, we wanted to elucidate how ATF4 becomes activated upon COX7a knock-down. ATF4 is known to be translated only under conditions of eIF2α phosphorylation, owing to the existence of two μORFs (open reading frames) in its 5’-UTR that block translation of the ATF4 ORF under normal conditions [22]. The *Drosophila* genome encodes two eIF2α kinases, GCN2 and PERK. GCN2 is activated by uncharged tRNAs, phosphorylating eIF2α to slow down translation when amino acids are lacking [22]. As the Dl^OE^, COX7a^RNAi^ proliferation was increased in an amino acid poor diet, the activation of GCN2 seemed the most likely explanation. However, we found that GCN2 knockdown did not rescue over-proliferation in by Dl^OE^, COX7a^RNAi^ animals, but induced higher larval and pupal lethality under nutrient restriction (Figure 4A). As these effects are largely independent of the genetic context, we assumed that GCN2-mediated translational repression via eIF2α phosphorylation was required for transient adaptation of the developing tissue to a lack of amino acids. The second eIF2α-kinase, PERK, is known for its role in mediating a transcriptional response through ATF4 as a result of ER protein folding stress [22,23], and interestingly, PERK knock-down completely rescued the over-proliferation in Dl^OE^, COX7a^RNAi^ animals (Figure 4A). This result suggested that PERK is activated upon COX7a knock-down, mediating eIF2α phosphorylation and consequently ATF4 translation. As expected, PERK^RNAi^ abolished the induction of ATF4 (Figure 4B), indicating that PERK acts upstream of ATF4.

**Figure 4:**
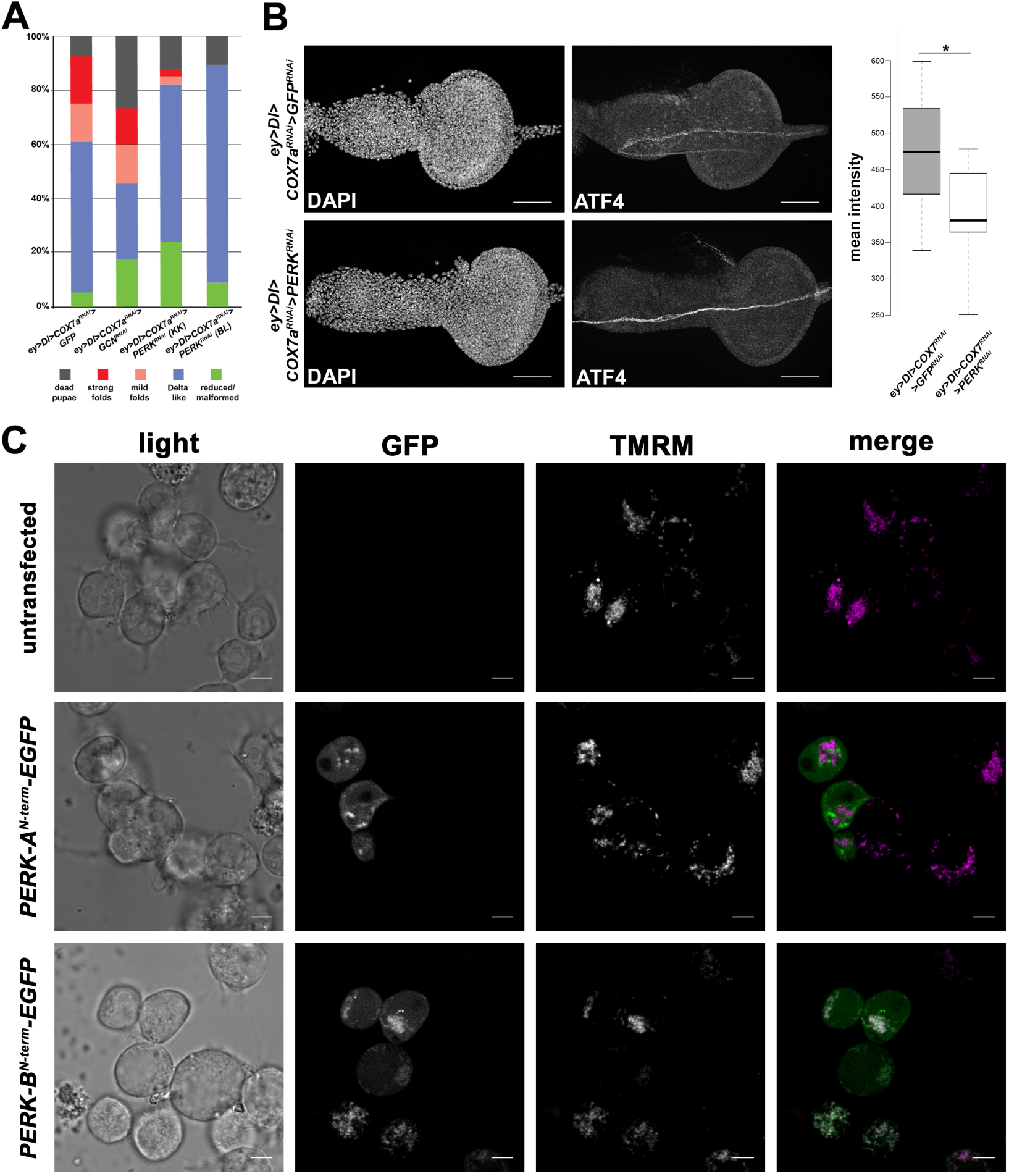
A mitochondria-targeted PERK isoform mediates ATF4 activation. **A** Quantification of adult eye phenotypes in crosses of *ey*>*Dl^OE^*,*COX7a^RNAi^* females with either UAS-GFP (control), *GCN2^RNAi^*, *ATF4^RNAi^* or two different *PERK^RNAi^* males. Larvae were reared at 25°C on nutrient restriction food. While knockdown of *GCN2* failed to abolish overgrown eyes, both *PERK^RNAI^* constructs repressed over-proliferation as did *ATF4^RNAi^*. **B** Max-projection of late L2 eye-antennal discs stained for DAPI and ATF4. Larvae were reared at 25°C on nutrient restriction food. Knock-down of PERK abolished any detectable ATF4 protein in Dl^OE^, COX7a^RNAi^ animals. Quantification (to the right) of GFP signal in early L3 eye discs of these genotypes. Asterisks indicate p-value according to unpaired t-test: ^*^ p<0,05; ^**^ p<0,01; ^***^ p<0,001. Anterior is to the left, dorsal is up, scale bar represents 50μm in microscope images. **C** Confocal single plane images of Drosophila S2R+ cells labelled with the mitochondrial dye TMRM. TMRM labelling highlights filamentous mitochondria in these cells. PERK-B^N-term^-EGFP is detected primarily in mitochondria, while PERK-A^N-term^-EGFP was found to be primarily excluded from mitochondria. Scale bar represents 5μm. See also Fig S4.

PERK is an ER-resident kinase that mediates one branch of the unfolded protein response of the endoplasmic reticulum (UPR^ER^) [23]. However, the ER chaperones BiP (Hsc70-3 in *Drosophila*) and GRP94 (Gp93 in Drosophila) as well as other well known targets of the Ire1/Xbp1 and ATF6 branches of the UPR^ER^ [24], were not induced according to our *in vivo* transcriptome profiling (Figure S4A). Consistently, the Xbp1-EGFP reporter [25] specific for this branch of the UPR^ER^ was not activated by COX7a knock-down at all (Figure S4B) and knock-down of ATF6 was not able to rescue the Dl^OE^, COX7a^RNAi^ induced over-proliferation (Figure S3E). Together, these results showed that the UPR^ER^ is not activated upon COX7a knockdown, but suggest a function of PERK independent of canonical ER stress.

The *Drosophila* genome encodes three isoforms of PERK, differing only in the N-terminus of the protein. Isoform A contains the expected N-terminal signal peptide for ER import, but we found isoform B to carry a potential mitochondrial import sequence according to MitoProtII prediction [26]. Isoform C contains no obvious signal peptide at all and is hardly expressed during development. To assess the potential mitochondrial localisation of isoform B, we fused the N-termini of PERK-A and PERK-B with EGFP and expressed these constructs in *Drosophila* S2R+ cells. As predicted, PERK-B^N-term^-EGFP showed a filamentous signal that overlapped with the mitochondrial dye TMRM (Figure 4C), while PERK-A^N-term^-EGFP showed a discrete labelling of the endomembrane system and did not co-localise with TMRM (Figure 4C).

Taken together, these results suggested that the PERK isoform B, which is targeted to mitochondria, acts as the eIF2α kinase activated upon COX7a knock-down, eliciting a transcriptional response through ATF4 that activated target genes of the Integrated Stress Response and the mitochondrial Unfolded Protein Response.

### ATF4-mediated adaptation is a general feature of ETC disturbance and enhances proliferation of other oncogenic pathways

Our results showed that ETC disturbance through COX7a knock-down activated the PERK-eIF2α-ATF4 axis, thereby inducing target genes of the ISR and UPR^mt^. As would be expected from a partial loss of ETC function, eye progenitors were compromised in their normal development despite the stress response, and these defects were further increased when the stress response was inhibited by ATF4^RNAi^ (Figure S3B). In contrast to this situation, activation of the Notch pathway by Dl^OE^ genetically interacted with ETC dysfunction to induce hyper-proliferation (Figures 1A-1B and 1D-1E), arguing that ATF4-mediated adaptation allows cells with elevated Notch signalling to further increase their rate of proliferation. To extend these findings beyond COX7a, we asked whether interference with other subunits of ETC complexes would cause similar phenotypes and ATF4 induction. In total we found 12 of 15 subunits (15/18 RNAi constructs) to cause similar or stronger phenotypes as COX7a^RNAi^ during eye development. All constructs targeting ETC Complex 1 or 4 induced an eye phenotype, while constructs targeting ETC Complex 2 showed no effects and only some constructs targeting Complex 3 resulted in eye phenotypes (Supplementary Table 2). The phenotypes induced were either very similar to COX7a^RNAi^ or showed severely reduced eyes and head malformations (Figure 5A), stronger than COX7a under enhanced knock-down conditions (Figure 1C and 1F). Importantly, we did not find any correlation of phenotypic classes with ETC complexes. To test for ATF4 induction, we used the LDH-GFP reporter and found that 3 of 3 Complex 1 subunits, 3 of 4 Complex 4 subunits and 1 of 2 Complex 3 subunits induced LDH-GFP expression (Figure 5B). Combined, these results strongly argued that knock-down of subunits of the large ETC Complexes 1, 3 and 4 induced an ATF4 response similar to COX7a^RNAi^. Though developmental phenotypes (reduced eyes or near-complete ablation of the eye primordium) and ATF4 induction are both common amongst the subunits we tested, the strength of these two phenotypes did not correlate well, indicating that cellular defects or stress might not necessarily be the signal activating ATF4 through PERK.

**Figure 5:**
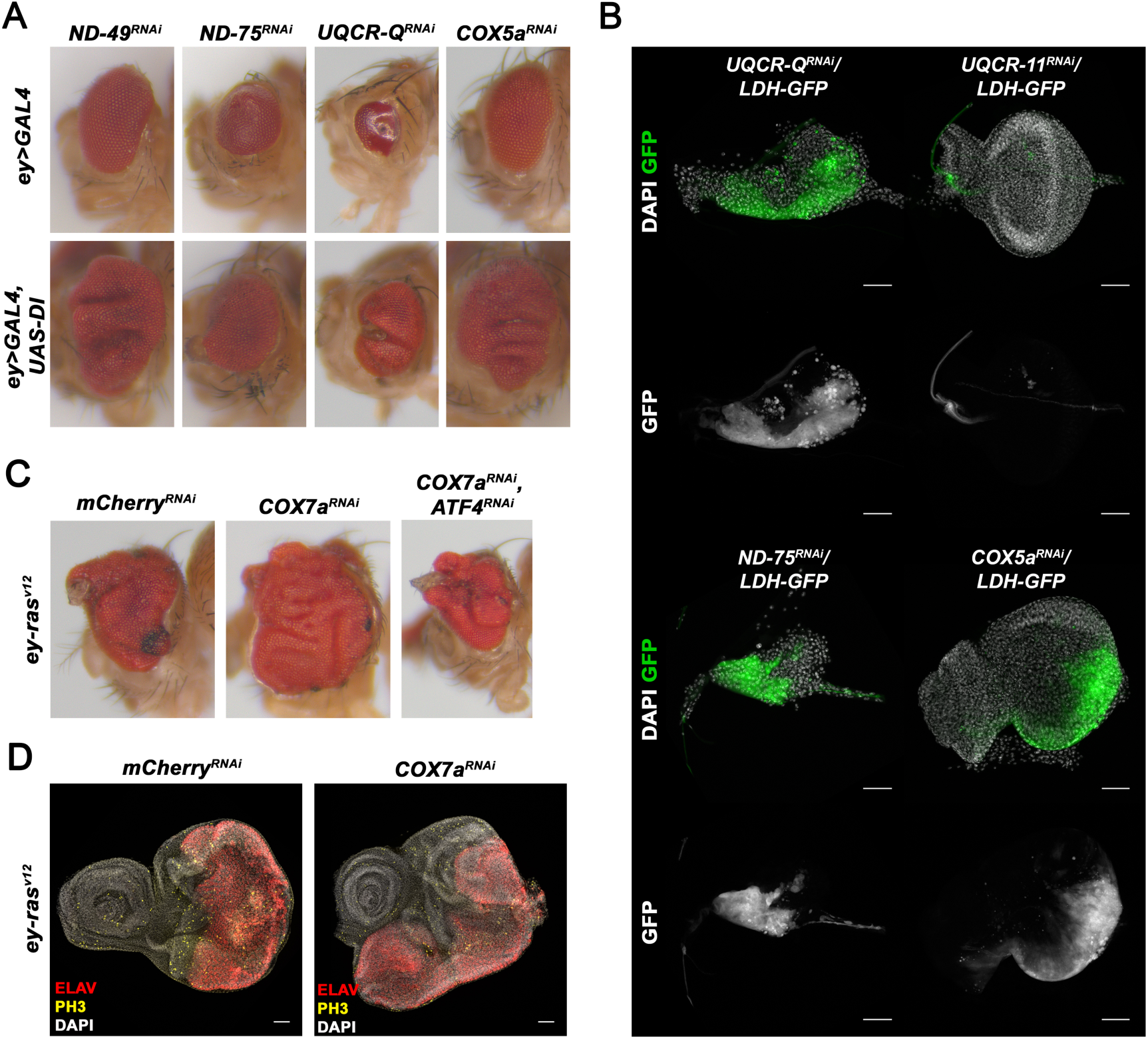
ATF4-mediated over-proliferation is a common cooperation of ETC knockdown and oncogenic signalling. **A** Adult eye phenotypes recovered upon knock-down of ETC subunits with either *ey*-GAL4 (upper row) or in the background of Dl^OE^ (lower row). ND-49 and COX5a caused slightly reduced eye size when individually knocked down (upper row) and over-proliferation with Dl^OE^. The strongly disturbed development in ND-75 or UQCR-Q knock-down eyes could not be transformed into over-proliferation by Dl^OE^. **B** Sum-projection of early L3 eye-antennal discs stained for DAPI (white, labelling nuclei) and showing GFP fluorescence of the *LDH*-*GFP* enhancer trap. The *LDH*-*GFP* reporter was induced upon knock-down of the indicated ETC subunits in varying strengths. Complex 3 subunit UQCR-11 did not induce GFP. **C** Adult eye phenotypes recovered upon COX7a knock-down in the background of ras^v12^ overexpression. In control knock-down (mCherry^RNAi^), eyes were slightly folded and often contained a characteristic dorsal-anterior outgrowth of eye and cuticle. COX7a^RNAi^ induced massive tissue folding, in an ATF4-dependent manner. **D** Sum-projection of late L3 eye-antenna discs stained for DAPI (white, labelling nuclei), ELAV (red, labelling photoreceptors) and phospho-Histone 3 (PH3) (yellow, labelling cells in mitosis). COX7a knock-down in the ras^v12^ over-expression background increased disc size and tissue folding. **E** Anterior is to the left, dorsal is up, scale bar represents 50μm in all microscope images. See also Fig S5.

After having established that knock-down of many subunits of different ETC complexes caused COX7a^RNAi^-like phenotypes, we sought to investigate whether the pro-proliferative genetic interaction with Notch signalling might be a more common feature as well. To this end, we used eye progenitors expressing oncogenic Ras^G12V^ (Ras^V12^), a common mutation in colon and lung carcinomas [27]. Ras^V12^ eyes are hyper-proliferative and show a characteristic outgrowth on the dorsal-anterior border of the eye as well as necrotic ommatidia (Figure 5C). Knock-down of COX7a dramatically increased eye size in age-matched larval discs (Figure 5D) and in adult survivors (Figure 5C and S5C). This effect was dependent on ATF4, as simultaneous interference with ATF4 reduced the over-proliferative phenotype (Figure 5C and S5C).

Together, these results demonstrated that reduced expression of ETC subunits caused an ATF4-mediated transcriptional response that in combination with growth-promoting signalling pathways like Notch or Ras leads to oncogenic over-proliferation of tissues.

## DISCUSSION

We showed here that genetically induced disturbance of ETC complexes resulted in a metabolic shift typical for mitochondrial impairment (Figures S5D-5F) [28,29] and activated an ATF4 dependent stress response. The *in vivo* transcriptional adaptation we present here confirmed the regulation of LDH and glycolytic enzymes as shown in *Drosophila* cultured cells [16] and further includes several targets shown to be ATF4 target genes in mammalian models [30,31]. Our results showed that the eIF2α-kinase PERK, so far only described for its role in mediating one branch of the UPR^ER^, is the upstream kinase phosphorylating eIF2α, thereby inducing ATF4 translation in response to mitochondrial ETC disturbance. Mitochondrial ETC^KD^ specifically activated PERK, while other branches of the UPR^ER^ are non-responsive. PERK activation upon mitochondrial defects was recently observed in *Drosophila* models of Parkinson’s disease [32] and was explained by the authors by its preferential localisation to mitochondria-associated ER membranes, which might make PERK more susceptible to a local stress signal. We have now shown here that *Drosophila* PERK isoform B contains a mitochondrial signal peptide, which is not found in mammalian PERK isoforms. The current evidence suggests that this *Drosophila* PERK isoform is imported into mitochondria, where it becomes degraded. Upon mitochondrial stress, import is disturbed and PERK accumulates in the cytoplasm, where it becomes activated. A similar strategy to monitor mitochondrial stress has been described in *C. elegans*, where an ATF4-like transcription factor is targeted to and degraded in mitochondria
[33].

In addition to canonical ATF4 target genes, we observed ATF4-dependent up-regulation of mitochondrial chaperones, a response classically referred to as the mitochondrial unfolded protein response (UPR^mt^). In *C. elegans*, mitochondrial chaperone induction upon stress is mediated by Atfs-1 [33], while the mammalian UPR^mt^ has been shown to be regulated by the evolutionary related transcription factor ATF5 [34]. Our data now showed that *Drosophila* ATF4 is required cell-autonomously for induction of mitochondrial chaperones upon ETC subunit knock-down, implying that *Drosophila* might represent the evolutionary ancestral ISR-UPR^mt^ regulation through a single ATF4-like transcription factor.

Previously, inhibition of mtDNA replication [35] and chemicals inhibiting different mitochondrial processes [31] have linked mitochondrial dysfunction to an ATF4 response. Our results presented here focus on the genetic knockdown of the nuclear-encoded ETC subunit 7a of Complex 4, but we have confirmed several other subunits of Complex 1, 3 and 4 to cause similar phenotypes and ATF4 induction *in vivo*, suggesting that genetic loss or reduction of individual subunits induces a common response. As upstream metabolic reactions did not elicit the same adaptation (data not shown), it appears unlikely that a general defect in metabolism would be the primary trigger of the ATF4 response. Therefore, our results are in agreement with the interpretation of Yoneda and colleagues [36], who suggested that the *C. elegans* UPR^mt^ is triggered in response to an overload of the mitochondrial protein folding machinery. This interpretation suggests that the ATF4 response is induced before detrimental dysfunction or damage occurs, possibly explaining why we failed to detect dysfunction or alterations of mitochondrial morphology (Figures S5A and S5B) in our model.

A novel finding presented in this study was the discovery that ATF4-mediated transcriptional adaptation allowed eye progenitors to increase their proliferation in response to signals from the Notch and Ras pathways. The primary questions arising from this genetic interaction is how these signalling pathways can overcome the apparent cellular stress and reduction in proliferation and then use the ATF4-mediated transcriptional adaptation for massively increased proliferation. A previous study showed that COX5a mutant clones activated a mitotic checkpoint through AMPK activation and reduced CycE expression [6]. Our cell culture metabolome studies suggested that AMP levels are indeed increased (Figure S5F). Furthermore, we observed a rescue of the COX7a knockdown induced small eye phenotype upon expression of CycE (data not shown), arguing that this phenotype might be primarily caused by a failure of cells to enter S-phase. As the Notch pathway had been shown to directly up-regulate CycE in *Drosophila* wing imaginal discs [37], Dl^OE^ in our model might simply overcome the checkpoint activated through rising AMP levels. In a second step, Notch signalling would be able to use the re-configured set of metabolic enzymes to drive cells faster through the cell cycle. Our transcriptome data imply that the differential capacity to take up nutrients from the hemolymph (such as amino acids) and the way they are used by the cells could be at the heart of this phenotype.

The cooperation between ATF4 target genes and the Notch or Ras pathways in *Drosophila* imaginal progenitors raised the intriguing possibility that these or other oncogenic pathways could benefit from ATF4 activity in human cancers. Over the last decades it had been demonstrated that human cancer cells are exposed to several stresses, including hypoxia, ROS or limitations in nutrient availability [38,39]. In order to survive these conditions and maintain their growth capacity, tumour cells activate responses like the HIF1α transcription factor axis [40]. Though less well studied, an involvement of ATF4 in cancer has been suggested mostly through work with cultured cells (for review see [41]). We analysed gene expression in human cancer samples of TCGA datasets using Cancer-RNAseq-Nexus [42] and the human protein pathology atlas [43], and found that many of the well characterised direct ATF4 targets [30] are up-regulated in a variety of cancer types. Most strikingly, transcriptomes of Kidney renal clear cell carcinoma showed progressive induction of ATF4 and many of its direct targets (EIF4EBP1, ASNS, TRIB3, VEGFA) on the transcriptional level, which strongly correlated with a poor prognosis in this type of cancer. In sum, our data suggest that the ATF4-mediated ISR is used by cancer cells to adapt their metabolic repertoire, thereby sustaining fast growth under increasingly unfavourable conditions.

## METHODS

### Fly stocks and husbandry

Flies were reared on a standard corn meal food recipe (cornmeal, barley malt, molasses, yeast extract, soy flour, propionic acid, Nipagin) at room temperature. Experimental crosses were carried out at 25°C unless otherwise noted. A reduced protein diet was prepared according to [8], with 20g fresh yeast per litre food and with 0.6% (v/v) propionic acid instead of Nipagin/Nipasol.

We used eyeless-Gal4 from [10] as the main Gal4-driver, but confirmed the primary phenotypes with the similar line ey3.5-Gal4 (BL-8220) and the unrelated driver so-Gal4 (BL-26810). RNAi lines were from Bloomington (TRiP libraries) or Vienna (KK library) stock centres. Whenever possible, phenotypes were confirmed by two independent RNAi lines (done for COX7a, COX5a, ND75). LDH-GFP was described in [14], TRE-GFP in [44] and Xbp1-EGFP in [25].

Rescue experiments in the ey>Delta,COX7aRNAi background were carried out with virgins from the triple transgenic stock (ey-Gal4, UAS-Delta, UAS-COX7aRNAi/CyO-TbRFP) crossed to males carrying UAS-RNAi or UAS-OE transgenes. As controls, UAS-GFP^nls^, UAS-mCD8-GFP, UAS-EGFP-RNAi or UAS-mCherry-RNAi were used. An average cross consisted of 10 to 12 virgins, mated with 6 to 10 males. Females were allowed to lay eggs for 1 to 2 days in fly food vials (10ml food on 5.3cm^2^ surface). To assess effects of crowding and for some crosses, freshly hatched L1 larvae were transferred by hand from apple-juice plates to food vials in defined numbers.

### Immunofluorescence

Antibody staining of larval eye-antennal discs was carried out according to standard procedures. In brief, larvae of corresponding stage (primarily early or late L3) were picked from the vial, collected, rinsed and dissected in PBS. Heads, usually consisting of mouth hooks, cephalopharyngeal skeleton, eye-antennal discs and CNS, were transferred to tube with PBS (with 0.01% Tween-20) on ice and then fixed with 4% para-formaldehyde for 18 minutes at room temperature on a nutator. Blocking was done with 1% blocking reagent (PerkinElmer) in PBT (0.1% Tween-20). Primary antibodies used were from DSHB (rat anti-ELAV 1:300, ms anti-Dachshund 1:300, ms anti-Wingless 1:40), Cell Signalling (rab anti-Casp3 1:200), Abcam (ms anti-ATP5a 1:500), SantaCruz (rab anti-PH3 1:1000) and Min-Ji Kang (gp anti-ATF4 1:66). Secondary antibodies were from Jackson Immunoresearch or Molecular Probes. Vectashield with DAPI was used as mounting medium.

### Microscopy and Image analysis

Larval discs were imaged on a Leica TCS SP8 confocal microscope. All experiments directly comparing antibody labelling of discs of different genotypes were dissected alongside, stained and imaged under identical settings (using master mixes for IF and identical laser and scanner settings). Images or confocal stacks were processed with the ImageJ distribution Fiji and Adobe Photoshop CS6. Generally, contrast was not enhanced. In some cases, rolling ball background subtraction was used. Most images shown are sum or maximum or sum projections of confocal stacks. Quantifications were done with Fiji: mean intensities were measured within rectangles centred at the equator of the eye field of sum projections or individual focal planes. Areas were manually outlined, taking folded parts of the tissue into consideration. Whole adult flies were photographed on a Zeiss Discovery V12 stereo microscope with a Zeiss HR3 camera.

### Transmission electron microscopy

Transmission electron microscopy (TEM) was carried out on late L3 larval eye-antennal discs. Eye-antennal discs were dissected in 1xPBS from third-instar larvae and fixed in 1.5% Paraformaldehyde and 0.2% Glutaraldehyde (GA) solved in 0.1 M Cacodylate buffer at 4°C over night. After four washing steps for 10 minutes with 0.1 M Cacodylate buffer the discs were fixed again with 2% Osmium in 0.1 M Cacodylate buffer for two hours at room temperature. Afterwards the discs were washed with 0.1 M Cacodylate buffer for 10 minutes twice and twice for 10 minutes with water (bidest). For raising the contrast of the samples the discs were incubated in 1% Uranyl acetate in water (bidest) at 4°C over night. After contrasting the discs were washed four times with water (bidest) and the water was subsequently stepwise replaced by ethanol. After five minutes of incubation in 100% Propylene oxide, the discs were incubated for an hour in 1:3 Epon/Propylene oxide, an hour in 1:1 Epon/Propylene oxide and finally replaced by pure Epon. For polymerisation the Epon was again replaced by fresh Epon and incubated for four hours at room temperature. The discs were transferred to beem capsules in Epon and the polymerization took place in an oven at 60°C. The samples were analysed with a Jeol JEM-1400 electron microscope (JEOL Deutschland, Freising) operated at 80 KV.

### RNA isolation and Microarray analysis

Total RNA was isolated from 20 to 30 early L3 eye-antennal discs using TRIzol (Invitrogen) and the Direct-Zol MicroPrep kit (ZymoResearch). 50ng of total RNA was used for processing with Affymetrix GeneChip^®^ 3’ IVT Express Reagent Kit (1st Microarray) or GeneChip^®^ 3’ IVT PLUS Reagent Kit (2nd Microarray). cRNA was hybridised to Affymetrix Drosophila Genome2.0 chips. The EMBL GeneCore Facility carried out IVT, hybridisation and chip imaging. Data was analysed with Affymetrix Software using the RMA normalisation algorithm. All fold-changes and significance (ANOVA p-value) reported in the study are based on this analysis.

### Analysis of metabolites

Free amino acids and thiols were extracted from 1^*^10^6^ S2R+ cells with 0.3 ml of 0.1 M HCl in an ultrasonic ice-bath for 10 min. The resulting extracts were centrifuged twice for 10 min at 4°C and 16.400 g to remove cell debris. Amino acids were derivatised with AccQ-Tag reagent (Waters) and determined as described in [45]. Total glutathione was quantified by reducing disulfides with DTT followed by thiol derivatisation with the fluorescent dye monobromobimane (Thiolyte, Calbiochem). For quantification of GSSG, free thiols were first blocked by NEM followed by DTT reduction and monobromobimane derivatisation. GSH equivalents were calculated by subtracting GSSG from total glutathione levels. Derivatisation was performed as described in [46]. UPLC-FLR analysis was carried out using an Acquity H-class UPLC system. Separation was achieved with a binary gradient of buffer A (100 mM potassium acetate, pH 5.3) and solvent B (acetonitrile) with the following gradient: 0 min 2.3 % buffer B; 0.99 min 2.3 %, 1 min 70 %, 1.45 min 70 %, and re-equilibration to 2.3 % B in 1.05 min at a flow rate of 0.85 ml min^-1^. The column (Acquity BEH Shield RP18 column, 50 mm × 2.1 mm, 1.7 μm, Waters) was maintained at 45°C and sample temperature was kept constant at 14 °C. Monobromobimane conjugates were detected by fluorescence at 480 nm after excitation at 380 nm and quantified using ultrapure standards (Sigma). Determination of organic acids was adapted from [47]. In brief, 1^*^10^6^ S2R+ cells per sample were extracted in 0.2 ml ice-cold methanol with sonication on ice. 50μl extract was mixed with 25μl 140 mM 3-Nitrophenylhydrazine hydrochloride (Sigma-Aldrich), 25 μl methanol and 100 μl 50 mM Ethyl-3-(3-dimethylaminopropyl) carbodiimide hydrochloride (Sigma-Aldrich) and incubated for 20 min at 60°C. Separation was carried out on the above described UPLC system coupled to a QDa mass detector (Waters) using an Acquity HSS T3 column (100 mm × 2.1 mm, 1.8 μm, Waters) which was heated to 40°C. Separation of derivates was achieved by increasing the concentration of 0.1 % formic acid in acetonitrile (B) in 0.1 % formic acid in water (A) at 0.55 ml min^-1^ as follows: 2 min 15% B, 2.01 min 31% B, 5 min 54% B, 5.01 min 90% B, hold for 2 min, and return to 15% B in 2 min. Mass signals for the following compounds were detected in single ion record (SIR) mode using negative detector polarity and 0.8 kV capillary voltage: Lactate (224.3 m/z; 25 V CV), malate (403.3 m/z; 25 V CV), succinate (387.3 m/z; 25 CV), fumarate (385.3 m/z; 30 V), citrate (443.3 m/z; 10 V), pyruvate (357.3 m/z; 15 V) and ketoglutarate (550.2 m/z; 25 CV). Data acquisition and processing was performed with the Empower3 software suite (Waters). The Metabolomics Core Technology Platform at the University of Heidelberg performed Metabolomic analysis.

### Cell culture

Drosophila S2R+ cells were cultured in Schneider’s medium (Gibco) with 10% FBS (Gibco). COX7a knockdown in S2R+ cells was done with *in vitro* transcribed dsRNA using a PCR template (targeting the identical sequence as the *in vivo* RNAi line). Cells were incubated with the dsRNA in serum-free medium for 45 minutes. Afterwards, serum concentration was adjusted to 10% again. Knockdown efficiency was monitored at 4 or 7 days after treatment by qPCR from TRIzol-extracted total RNA. For transient transfections, Qiagen Effectene Transfection Reagent was used. Constructs were constitutively expressed from an actin promoter (pAC5.1_DEST, gift of M. Boutros). Live-imaging was carried out one or two days post-transfection in glass-bottom dishes (MatTek).

## AUTHOR CONTRIBUTIONS

S.S. conceived the study, designed, performed and interpreted most experiments, supervised and worked with J.T. und C.A. and wrote the paper with I.L. J.T. performed and interpreted experiments on COX7a knockdown in cultured S2R+ cells and performed experiments on food dependency of COX7a phenotypes. C.A. assisted in a primary RNAi screen, designed and performed the TEM experiment and helped with initial experiments. I.L. conceived the study, assisted in designing and interpreting experiments, wrote the paper with S.S., obtained funding to support the study DFG (Excellence Cluster “CellNetworks/EcTop6). S.S. obtained a PhD fellowship from HBIGS.

## ACKNOWLEDGMENTS

We thank Jelena Pistolic and Vladimir Benes of EMBL GeneCore for the Microarray analysis. We thank the Metabolomics Core Technology Platform of the Excellence cluster “CellNetworks” (University of Heidelberg) and the Deutsche Forschungsgemeinschaft (grant ZUK 40/2010-3009262) for support with HPLC-based metabolite quantification. We thank Steffi Gold, as well as Stefan Hillmer at the Heidelberg University Electron Microscopy Core Facility (EMCF), for their help with the EM component of the work. Finally, we are indebted to Min Ji Kang, Stefanie Schirrmeier, Michael Boutros, Aurelio Teleman and Ana Gomez Barata for fly lines or reagents. We would also like to thank A. Teleman, J. Friedrich, P. Pinto and K. Domsch for carefully reading the manuscript and N. Trost for help with the GO term analysis. This work was supported by the Excellence Cluster CellNetworks (EcTop6) and a HBIGS PhD fellowship to S.S. We apologize to all whose work was not cited due to space limitations.

